# Hormonally responsive bovine oviductal organoids recapitulate native oviductal secretions and enhance sperm capacitation

**DOI:** 10.64898/2026.03.10.710777

**Authors:** Sergio Navarro-Serna, Jon Romero-Aguirregomezcorta, Nathaly Hernández-Díaz, Ana Ferrero-Micó, Pilar Coy, Vicente Pérez-García

## Abstract

The oviduct provides the dynamic microenvironment that supports fertilization and early embryo development yet replicating its hormonally regulated secretory activity *in vitro* remains a major challenge. Here, we established bovine oviductal epithelial organoids that reproduce the structural polarity and endocrine responsiveness of the native oviduct. Exposure to either estradiol or progesterone resulted in distinct transcriptomic and proteomic landscapes that were characteristic of the follicular and luteal phases, respectively. This included the upregulation of canonical phase-specific markers, such as *OVGP1*, *NTS*, *HP* and *TGM2*. Proteomic profiling of organoid-derived secretions (ODS) revealed extensive overlap with *in vivo* oviductal fluid. Integration of transcriptomic and proteomic datasets by multi-omics factor analysis identified coherent biological signatures defining each hormonal state. Functionally, ODS obtained from estradiol-treated organoids enhanced sperm capacitation and acrosome reaction, recapitulating the activity of follicular-phase oviductal fluid. These findings demonstrate that hormonally responsive oviductal organoids generate bioactive secretions that emulate the molecular and functional features of the native oviductal environment, providing a sustainable and physiologically relevant platform for studying gamete-maternal communication and improving assisted reproduction technologies.

## Introduction

Fertilization and early embryonic development in mammals occur within the oviduct, a highly specialized segment of the female reproductive tract that provides the dynamic biochemical and biophysical cues required for gamete maturation, fertilization, and preimplantation embryo development [1, 2]. The oviductal epithelium orchestrates this environment through hormonally regulated secretory activity, producing a complex mixture of proteins, lipids, metabolites, and extracellular vesicles that interact with both gametes and embryos [3, 4]. This finely tuned secretory function fluctuates across the oestrous cycle, driven by estradiol and progesterone, which regulate epithelial differentiation, ciliary activity, and molecular transport [4]. Disruptions to this signalling or to the composition of oviductal secretions can impair sperm capacitation, fertilization efficiency, and embryo quality [5, 6].

Despite extensive research on the composition of oviductal fluid, replicating its dynamic and hormone-dependent physiology *in vitro* remains a major challenge. Traditional two-dimensional cultures of bovine oviductal epithelial cells have provided valuable insights into hormone responsiveness and gamete interactions [7] but lack the epithelial polarity and luminal compartmentalisation necessary to mimic *in vivo* secretion. Direct use of oviductal fluid from slaughterhouse material has demonstrated beneficial effects on sperm function by modulating sperm capacitation [8, 9] and on embryo quality by increasing post-thaw survival and modulating embryo epigenetic patterns [10–12]; however, variability among donors, and degradation of secretions after tissue collection restrict its reproducibility and translational value.

Three-dimensional (3D) culture systems offer new opportunities to model the oviductal microenvironment under physiologically relevant conditions [13]. Studies using other 3D models of the bovine oviduct, such as spheroids, have reported successful strategies to study embryo–maternal interactions [14–16]. The apical-out orientation allows the investigation of physical interactions with gametes and embryos; however, this model presents important limitations, including the inability to be passaged and the fast loss of their secretory profile within a few days [14]. Consequently, while spheroids are suitable for studying physical interactions, they are not optimal models for investigating embryo–maternal paracrine communication. Organoids derived from reproductive epithelia self-organize into polarized structures that recapitulate key morphological and functional features of the native tissue [17, 18] The apical-in organization of organoids allows the study and isolation of luminal secretions produced by the epithelium without contamination from culture medium components, as previously demonstrated. [19]. In particular, uterine epithelial organoids maintain long-term expandability and hormone responsiveness and secrete biomolecules into their enclosed lumen, creating a controlled *in vitro* source of uterine secretions [18, 20]. Studies in human and murine endometrial organoids have shown that endocrine stimulation can reproduce phase-specific transcriptomic profiles [18, 20], however, this strategy has yet to be fully explored in the bovine oviduct. The bovine species represents a valuable reproductive model, as its pre-implantation embryo development closely resembles that of humans in terms of timing, metabolic activity, and molecular regulation, providing a relevant system for translational reproductive research [21].

Here, we report the establishment of hormonally responsive bovine oviductal epithelial organoids that reproduce the transcriptional and functional signatures of the oestrous cycle. By integrating transcriptomic and proteomic analyses, we demonstrate that estradiol and progesterone treatments elicit distinct molecular programs characteristic of the follicular and luteal phases, respectively. The organoid-derived secretions (ODS) closely recapitulate the composition of native oviductal fluid and, importantly, functionally enhance sperm capacitation and acrosome reaction. This study provides a mechanistic framework for modelling oviductal physiology *in vitro* and introduces a renewable, ethically sustainable platform to study gamete–maternal interactions and to refine assisted reproductive technologies.

## Material and methods

### Organoid derivation and culture

Bovine reproductive tracts were obtained from commercial cattle at a slaughterhouse and processed following a modified protocol based on Romar et al. (2001) [22]. The tracts were washed once in 0.04% cetrimide solution and twice in physiological saline solution. Oviducts were then clamped with sterile forceps and rinsed in phosphate-buffered saline (PBS) supplemented with 10% penicillin/streptomycin. To isolate epithelial cells, the clamped oviducts were filled with Trypsin-EDTA solution and incubated at 38.5 °C for 45 minutes. Cells were subsequently collected by a second flush with Trypsin-EDTA solution, and the entire content was transferred into a conical tube containing RPMI 1640 medium (Thermo Fisher Scientific, 11875093) supplemented with 10 % (v/v) fetal bovine serum (FBS), 50 IU penicillin, 50 µg/mL streptomycin and 2.5 µg/mL amphotericin B (Sigma-Aldrich, A2942) After centrifugation at 800 × g for 4 minutes, the supernatant was discarded, and the pellet was resuspended in CryoStor CS10 (Sigma-Aldrich, C2874) and stored in cryovials at -80 °C until further processing.

Organoid derivation procedure was based on Turco et al. (2017) [18] with minor modifications. Samples were thawed and washed with RPMI 1640 medium. After centrifugation at 500 × g for 5 minutes, the pellet was resuspended in a digestion solution (1 mg/mL collagenase V, Sigma, C-9263; 0.1 mg/mL DNase I, Roche, 1128493, diluted in RPMI 1640 supplemented with 10% FBS) and incubated at 37 °C with shaking for 30 minutes. Following centrifugation under the same conditions, the pellet was resuspended in PBS, passed through a 40 µm sieve, and the sieve was washed twice with PBS. The sample was centrifuged again, and the pellet was resuspended in Advanced DMEM/F12 (Gibco, 12634010) supplemented with 10% FBS and 5% penicillin/streptomycin.

To isolate fibroblasts, samples were seeded into Petri dishes and incubated for 1 hour. After incubation, the medium containing non-adherent cells was collected, and fresh medium was added to the Petri dish. The suspension of non-adherent cells was centrifuged under the same conditions, washed with Advanced DMEM/F12, and centrifuged again. The resulting pellet was resuspended in Matrigel (Corning, 356231), and 25 µL drops were plated into 48-well plates. After solidification, each drop was overlaid with 250 µL of Expansion Medium as described by Turco et al. (2017). The medium was refreshed every 2–3 days.

For passaging, the organoid droplets were collected in a 1.5 mL tube and mechanically dissociated by pipetting 200 times in two separate steps, with a wash in Advanced DMEM/F12 between them. For storage, the same procedure was followed but with only one dissociation step; after washing, the pellet was resuspended in CryoStor CS10.

### Hormonal stimulation

To mimic the hormonal profiles of the follicular and luteal phases, organoids were treated for 4 days with two hormone treatments adapted from the endometrial organoid differentiation protocol described by Turco et al. (2017) [18], with modifications for the bovine species. For follicular phase mimicking, organoids were treated with β-estradiol (E2, Sigma, E4389) at 10 nM in Expansion Medium for 4 days. For luteal phase mimicking, organoids were first treated with 10 nM E2 for 2 days, followed by an addition of 2 days with 1 nM E2, 1 µM medroxyprogesterone acetate (mP4, Sigma, PHR1589), and 1 µM 8-bromoadenosine 3′,5′-cyclic monophosphate (cAMP, Sigma, B7880). A control group without hormonal treatment was also included, with the same concentration of hormonal extenders as in the experimental groups.

### Organoid derived secretion and organoid pellet collection

ODS were extracted as described by Simintiras et al. 2021 [23]. Matrigel was removed using Cell Recovery Solution (Corning, 3254253), and the resulting organoid suspensions were washed in PBS to eliminate residual Cell Recovery Solution. Samples were centrifuged at 300 × g for 10 minutes at 4 °C. The pellets were resuspended in PBS, and organoids were disrupted first by pipetting (100 times) and subsequently by vortexing for 5 minutes. Samples were then centrifuged at 3750 × g for 15 minutes at 4 °C. The supernatant, containing ODS diluted in PBS, was collected, while the cellular pellets were retained for RNA or protein extraction.

### Sperm capacitation treatment

Frozen semen straws from five different Asturiana de los Valles bulls (SERIDA, Deva, Spain), proven for both artificial insemination and *in vitro* fertility, were used in the experiments. For each replicate, at least three 0.25 mL straws were thawed per bull at 38°C for 30 seconds and pooled to minimize inter-straw variability. The samples were then diluted in Sperm Wash Medium (EmbryoCloud, Murcia, Spain; 19990/0130) and centrifuged at 700 × g for 3 minutes. The resulting sperm pellet was resuspended in Fertilization Medium (EmbryoCloud; 19990/0140) to achieve a final concentration of 20 million spermatozoa per mL. Following the different experimental conditions, spermatozoa were incubated for 2 hours at 38 °C in 5% CO_2_. After sperm capacitation, five samples were used to evaluate motility and membrane status, and four samples were centrifuged for 5 minutes at 1000 × g; the resulting pellets were prepared for western blot analysis.

### RNA extraction and RT-qPCR

Total RNA was extracted using the RNeasy Mini Kit with RNase-free DNase treatment (Quiagen, 74106). Retrotranscription was performed to synthesized complementary DNA using PrimeScript RT Master Mix (Takara Bio, RR036A). Quantitative real-time PCR was performed with TB Green Premix Ex Taq (Takara Bio, RR420L) on a LightCycler 480 II thermocycler using 348-well plates. Gene expression was normalized to the geometric mean of GAPDH and actin, which served as housekeeping genes. The complete list of primers is provided in Supplementary Table 1. For statistical analysis, data are presented as mean ± SEM and differences in gene expression were assessed using one-way ANOVA followed by Tukey’s multiple comparison test (p≤0.05) in GraphPad Prism 9.

### Western Blot

Samples for Western blot analysis, including both organoid and sperm pellets, were lysed in RIPA buffer (50 mM Tris-HCl, 150 mM NaCl, IGEPAL CA-630, 0.75% sodium deoxycholate, 0.1% SDS, and protease inhibitors) and denatured in loading buffer by heating at 95 °C for 5 minutes. Proteins were separated on 8% SDS-PAGE gels using Precision Plus Protein Dual Color Standards (Bio-Rad) as molecular weight markers and transferred onto PVDF membranes (Millipore). Membranes were blocked with 5% milk in TBS-T (1× Tris-buffered saline with 0.1% Tween-20) or 5% bovine serum albumin (BSA) in TBS-T when pPKA and pTyr were incubated.

Blots were incubated with primary antibodies against TGM2 (1:1000, Invitrogen, MA5-32819), Phospho-PKA Substrate (1:1000, Cell Signaling, 9624), p-TYR (1:1000, Sigma-Aldrich, 4G10), β-Tubulin (1:5000, Sigma-Aldrich, T0198) and Actin (1:5000, Abcam, ab6276). After washing, membranes were incubated with horseradish peroxidase-conjugated secondary antibodies anti-rabbit (1:3000, Bio-Rad, 170–6515) and anti-mouse (1:3000, Bio-Rad, 170–6516). Detection was performed using an enhanced chemiluminescence kit (GE Healthcare, RPN2209) and visualized on X-ray films. Band intensities were quantified with ImageJ software. For statistical analysis, data are presented as mean ± SEM and differences in relative intensity were assessed using one-way ANOVA followed by Tukey’s multiple comparison test (p≤0.05) in GraphPad Prism 9.

### Immunofluorescence staining

Organoids were released from Matrigel using Cell Recovery Solution (Corning, 3254253) and washed with PBS to remove residual matrix. The suspensions were centrifuged at 300 × g for 10 min at 4 °C, and the resulting pellets were fixed in 4% paraformaldehyde in PBS for 15 min at room temperature. After fixation, organoids were permeabilized with 0.5% Triton X-100 and 0.1% BSA in PBS for 3 h at RT. Following three PBS washes (15 min), organoids were blocked overnight at 4 °C in the blocking buffer (permeabilization buffer supplemented with 2% FBS). Samples were then incubated overnight at 4 °C with primary antibodies against ESR2 (1:100, Abcam, ab5786), CDH1 (1:100, BD Biosciences, 610181), acetyl-α-tubulin (1:100, Cell signaling, 5335) and SOX17 (1:100, R&D System, AF1924) diluted in blocking buffer. After three washes (15 min) in blocking buffer, organoids were incubated overnight at 4 °C with Alexa Fluor 488 or Alexa Fluor 568–conjugated secondary antibodies (1:400, Thermo Fisher Scientific). Subsequently, organoids were washed three times for 1 h in blocking buffer, and nuclei were counterstained with DAPI for 10 min at RT. Finally, organoids were mounted in Vectashield mounting medium (Vector Laboratories, H-1000) within a support frame to prevent compression. Images were acquired using an LSM900 vertical confocal microscope (Zeiss).

### Flow cytometry

Sperm viability and acrosome integrity were assessed by flow cytometry using propidium iodide (PI) and fluorescein isothiocyanate-labeled peanut agglutinin (FITC-PNA) on a Guava easyCyte® system (Merck Millipore, Madrid, Spain) with standard FL1/FL3 detectors, as previously described [24]. Sperm were diluted to 1–2 × 10⁵ cells/mL in a staining solution containing PI (500 µg/mL) and FITC-PNA (200 µg/mL) in PBS and incubated for 15 min at room temperature in the dark. Fluorescence compensation corrected spectral overlap. Sperm were classified as: viable with intact acrosome (PI⁻/FITC-PNA⁻), viable with damaged acrosome (PI⁻/FITC-PNA⁺), non-viable with damaged acrosome (PI⁺/FITC-PNA⁺), or non-viable with intact acrosome (PI⁺/FITC-PNA⁻). Membrane fluidity was evaluated by co-staining with merocyanine 540 (M540) and YoPro-1. Sperm were incubated with Merocyanine 540 (M540, 2.6 µM) and YoPro-1 (25 nM) for 10 min at 37 °C in the dark. Fluorescence was detected using FL3 (M540) and FL1 (YoPro-1) channels. Sperm were classified into four populations: viable with low membrane lipid disorder (M540⁻/YoPro-1⁻), viable with high membrane lipid disorder (M540⁺/YoPro-1⁻), non-viable with low membrane lipid disorder (M540⁻/YoPro-1⁺), and non-viable with high membrane lipid disorder (M540⁺/YoPro-1⁺). For statistical analysis, data are presented as mean ± SEM and percentage of spermatozoa with high membrane fluidity, acrosome reaction and viability were assessed using one-way ANOVA followed by Tukey’s multiple comparison test (p≤0.05) in GraphPad Prism 9.

### Analysis of sperm motility

Sperm motility was assessed using a computer-assisted sperm analysis (CASA) system (ISAS V1, Proiser, Valencia, Spain). Chambers (20 µm depth; Spermtrack, Proiser) were pre-warmed to 38 °C. A 4 µL drop of each seminal sample was loaded, and at least three fields totaling >200 spermatozoa per sample were analysed.

The CASA parameters measured included progressive motility (%), curvilinear velocity (VCL, µm/s), straight-line velocity (VSL, µm/s), average path velocity (VAP, µm/s), linearity (LIN = VSL/VCL, %), straightness (STR = VSL/VAP, %), wobble (WOB = VAP/VCL, %), amplitude of lateral head displacement (ALH, µm), and beat cross frequency (BCF, Hz). Analysis settings were 100 frames per field, with sperm tracked for at least 15 frames and 2 s at 50 frames/s. Imaging was performed using a negative-contrast phase microscope (100×; Leica DMR, Wetzlar, Germany) coupled to a digital camera (Basler Vision, Ahrensburg, Germany). For statistical analysis, data are presented as mean ± SEM and data were assessed using one-way ANOVA followed by Tukey’s multiple comparison test (p≤0.05) in GraphPad Prism 9.

### RNA-seq sample preparation

Organoids RNA samples were extracted using RNeasy mini-Kit and RNase-free DNase (QIAGEN 74106). QC, library preparation, sequencing, and mapping were performed by Novogene using an Illumina Novaseq 6000 with 150bp paired-end reads. Sequence data were deposited in ENA under accession PRJEB104075.

### Proteomic LC-MS/MS

Proteomic analysis of ODS was performed using liquid chromatography–tandem mass spectrometry (LC–MS/MS) at the Proteomics Service of the Servicio Central de Soporte a la Investigación Experimental, University of Valencia. ODS recovered in PBS were quantified, and 10 µg of total protein per sample was used for analysis.

Sample preparation followed the single-pot, solid-phase–enhanced sample preparation (SP3) protocol for detergent removal and clean-up prior to enzymatic digestion with trypsin, as described by Moggridge et al. (2018) and Müller et al. (2020) [25, 26] with minor modifications. Following digestion, 2 µL of each sample were diluted to a final volume of 20 µL with 0.1% formic acid and loaded onto Evotip Pure tips (EvoSep) according to the manufacturer’s instructions.

Peptide identification was performed using MSFragger (via FragPipe) against the SwissProt human protein database. Protein quantification and differential expression analyses were carried out using the default DIA-NN workflow. The mass spectrometry proteomics data have been deposited to the ProteomeXchange Consortium via the PRIDE partner repository with the dataset identifier PXD071140.

### Bioinformatics analysis

RNA-seq data were processed and quantified by Novogene to generate a gene-level count matrix. Downstream analyses were performed in R (v4.3.1) using Bioconductor and tidyverse packages. Raw counts were batch-corrected using ComBat_seq (sva v3.46.0) and normalized to TPM values. Differential gene expression was assessed with DESeq2 (v1.42.0) applying Wald tests and Benjamini–Hochberg false discovery rate (FDR) correction. Genes with p < 0.05 and |log₂ fold change| > log₂(1.5) were considered significantly differentially expressed. Principal component analysis (PCA) and volcano plots were generated using ggplot2, and Gene Ontology enrichment analysis was subsequently performed.

The cellular composition of bovine oviductal organoids was estimated via RNA-seq deconvolution using a signature matrix derived from mouse single-cell RNA-seq (scRNA-seq) data from sample GSM5005819 [27]. The scRNA-seq dataset was processed with Seurat [28]. Top 25 markers genes previously defined for each cell type in McGlade et al. 2021 were defined, and modulate scores were calculated to assign predominant cell types to each cluster. The mouse signature matrix was mapped to bovine genes and the bulk bovine RNA-seq matrix was filtered to match the signature genes. Deconvolution was performed using CIBERSORTx [29] and the estimated cell fraction per sample were visualized with bar plots.

Label-free quantitative (LFQ) proteomics data were analysed in R following a workflow based on the DEP package (v1.26.0) combined with Bioconductor and tidyverse utilities. Proteins missing in any sample were excluded, and remaining intensities were normalized by variance stabilizing normalization (VSN). Batch effects were corrected using limma’s removeBatchEffect function, including experimental condition as a covariate. Proteins with p < 0.05 and |log₂ fold change| > log₂(1.5) were considered differentially regulated. Visualization of results was performed using ggplot2 and ggrepel.

For integrative analysis, log₂-transformed TPM transcriptomic data and batch-corrected LFQ proteomic intensities were combined using MOFA2 to identify latent factors capturing shared and modality-specific sources of variation. Low-variance features were filtered out prior to model training. Factor scores were visualized by experimental group and batch, and the top contributing genes and proteins for selected factors were extracted for downstream functional interpretation.

## Results

### Generation and hormonal responsiveness of bovine oviductal epithelial organoids

Bovine epithelial organoids were established from epithelial cells isolated from the infundibular–ampullar region of slaughterhouse-derived reproductive tracts (Fig. 1A). When embedded in Matrigel, cells self-organized into cystic 3D structures composed of a single epithelial monolayer (Fig. 1B). Immunofluorescence staining confirmed the presence of the epithelial marker E-cadherin (CDH1) outlining the organoid surface and a laminin-positive basal membrane (Fig. 1C), consistent with apical-in polarization typical of oviductal organoids.

**Figure 1.**
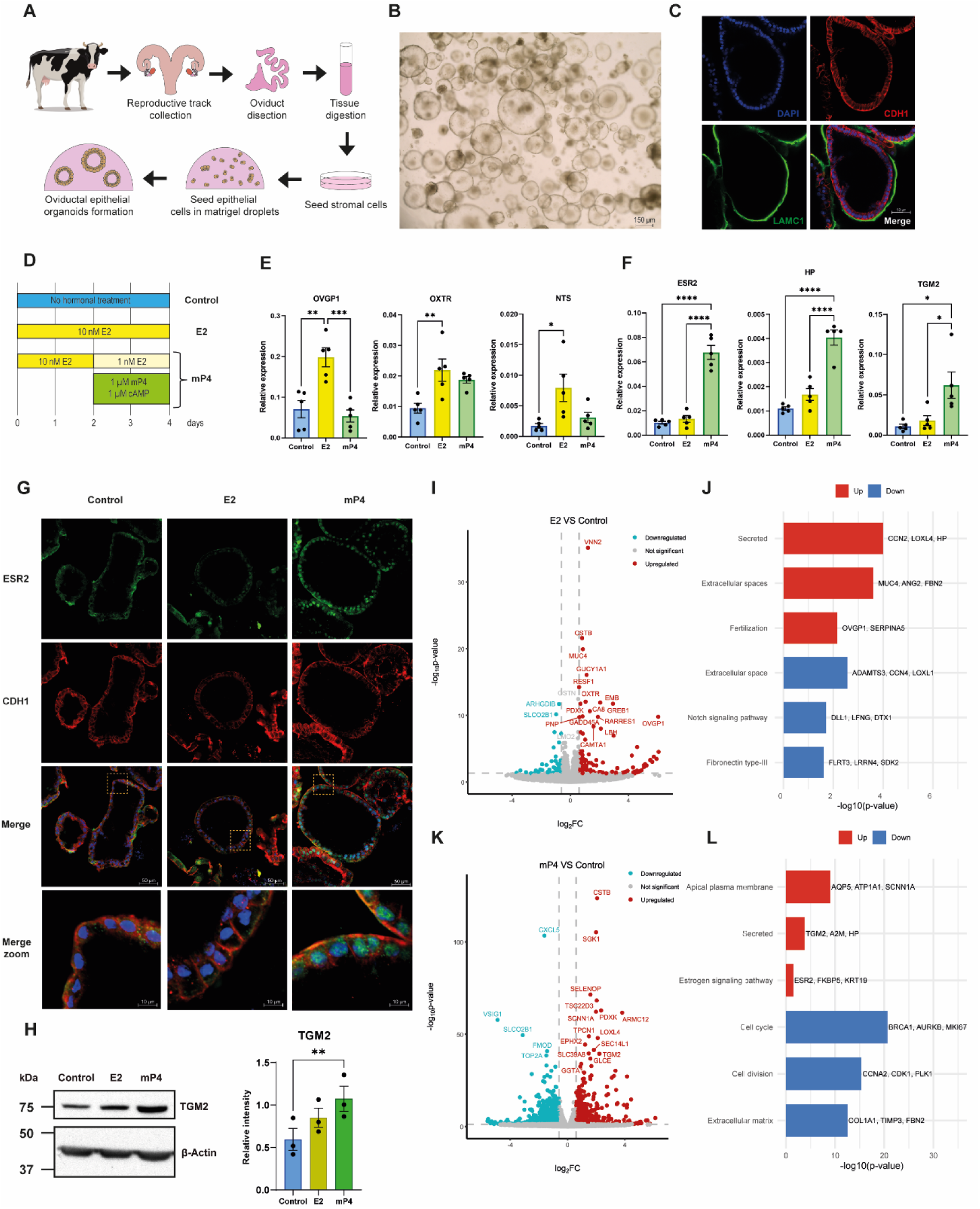
Generation of bovine oviductal epithelial organoids and characterization of oestrous cycle hormone mimicking. (A) Schematic representation of the generation of bovine oviductal epithelial organoids. (B) Bright-field image of bovine oviductal epithelial organoids before passage. (C) Immunofluorescence staining of organoids after hormonal treatments with Laminin (green, basal layer marker), E-cadherin (red), and DAPI (blue). (D) Hormonal treatment protocol. Control: no hormonal treatment, with the addition of hormone vehicle; E2: 10 nM estradiol for 4 days; mP4: 10 nM estradiol for 2 days followed by 1 nM estradiol, 1 µM medroxyprogesterone acetate (mP4), and 1 µM 8-bromoadenosine 3′,5′-cyclic monophosphate sodium salt (cAMP). (D–E) Gene expression analysis by RT-qPCR of markers of the follicular (E) and luteal (F) phases, n = 5 independent biological replicates. (G) Immunofluorescence staining of organoids after hormonal treatments with ESR2 (green, luteal-phase marker), E-cadherin (red), and DAPI (blue). (H) Western blot of TGM2 (luteal-phase marker) upregulated in mP4 group and Actin such as housekeeping and signaling quantification relative to the housekeeping, n = 3 independent biological replicates. (I) Volcano plot of differentially expressed genes in the E2 group vs. Control, n = 3 independent biological replicates. (J) Gene ontology enrichment of differentially expressed genes between E2 and Control groups; terms upregulated by estradiol treatment are shown in red, and downregulated terms in blue. (K) Volcano plot of differentially expressed genes in the mP4 group vs. Control, n = 3 independent biological replicates. (L) Gene ontology enrichment of differentially expressed genes between mP4 and Control groups; terms upregulated by mP4 treatment are shown in red, and downregulated terms in blue. For statistical analysis of RT-qPCR and western blot quantification, differences were assessed using one-way ANOVA followed by Tukey’s multiple comparison test (p ≤ 0.05). Data are presented as mean ± SEM. Statistical significance is indicated as *p < 0.05, **p < 0.01, and ***p < 0.001.

To reproduce the endocrine dynamics of the oestrous cycle, organoids were exposed to estradiol (E2) or to a combination of estradiol, progesterone analog (mP4), and cAMP (Fig. 1D). RT–qPCR revealed a phase-specific transcriptional response: E2 stimulation upregulated *OVGP1*, *NTS*, and *OXTR*, canonical follicular-phase markers, whereas mP4 treatment increased expression of luteal-phase markers including *HP*, *TGM2*, and *ESR2* (Fig. 1E–F). Hormonal regulation of selected markers was further validated by immunofluorescence for ESR2, which showed enhanced nuclear localization in the mP4-treated group (Fig. 1G), and by western blot, confirming TGM2 overexpression in organoids exposed to mP4 (Fig. 1H). Together, these results demonstrate that bovine oviductal organoids preserve epithelial integrity and exhibit steroid hormone responsiveness, recapitulating the transcriptional and functional dynamics of the *in vivo* oestrous cycle.

### Estradiol and progesterone treatments elicit distinct transcriptomic signatures reflecting follicular and luteal phase physiology

To define the molecular basis of hormone responsiveness, we performed RNA-seq on organoids cultured under control, E2, and mP4 conditions. As shown in Supplementary Figure 1A, the control and estradiol-treated groups exhibited similar profiles, in contrast to the progesterone-treated group. Compared to the control, estradiol treatment resulted in 225 differentially expressed genes (DEGs; 155 upregulated and 70 downregulated; p ≤ 0.05, |log₂ fold change| > log₂(1.5)), whereas mP4 treatment affected 1,330 genes (463 upregulated and 867 downregulated); Fig. 1I, K and Supplementary Table 2). The transcriptomic responses closely recapitulated *in vivo* patterns reported for the bovine oviduct. Specifically, comparison with the microarray dataset from Cerny et al. (2015) [30] revealed substantial overlap, with 142 follicular-phase and 107 luteal-phase genes showing concordant upregulation in our model (Supplementary Table 3). These findings demonstrate that the organoid system effectively reproduces phase-specific transcriptional programs characteristic of the oestrous cycle.

Gene Ontology (GO) enrichment analysis of E2-upregulated genes revealed activation of pathways associated with secretion, extracellular remodelling, and oviductal luminal composition, including *OVGP1*, *SERPINA5*, and *MUC4*. Downregulated terms were enriched in Notch signalling and epithelial differentiation, suggesting estradiol promotes a secretory phenotype while transiently repressing differentiation cues (Fig. 1J, Supplementary Table 4). Progesterone treatment, in contrast, enhanced the expression of genes linked to secreted proteins characteristic of luteal phase (*TGM2, HP*), apical plasma membrane remodelling (*AQP5*, *ATP1A1*), and metabolic adaptation, while repressing cell-cycle regulators (*BRCA1*, *CDK1*) and extracellular matrix components (*COL1A1, FBN2*) (Fig. 1L, Supplementary Table 4). Deconvolution of bulk RNA-seq (Supplementary Figure 1B) further indicated that ampullar secretory cells constitute the main cell population in the organoids. This was confirmed by immunofluorescence, showing uniform SOX17 expression (a marker of secretory cells, Supplementary Figure 1C) and the absence of multiciliated cells, as visualized by primary cilia staining (Supplementary Figure 1D). No differences were observed between hormonal treatment groups in either staining. Collectively, these data indicate that estradiol promotes a secretory and proliferative state, whereas progesterone drives epithelial maturation, reduced proliferation, and extracellular stabilization, mirroring the transition from follicular to luteal physiology.

### Hormonal stimulation mimics the proteomic composition of organoid-derived secretions

Given that the oviductal epithelium exerts its reproductive functions largely through its secretions, we next characterized the protein content of ODS (Fig. 2A). Liquid chromatography–tandem mass spectrometry identified 4,070 proteins across all conditions, many of which overlap with previously reported bovine oviductal fluid proteomes. A qualitative comparison with *in vivo* oviductal fluid collected during the early luteal phase further revealed that ODS shared 83.7–83.9% of their protein content with native oviductal fluid, underscoring their capacity to faithfully reproduce the physiological oviductal environment (Supplementary Figure 2 A-C).

**Figure 2.**
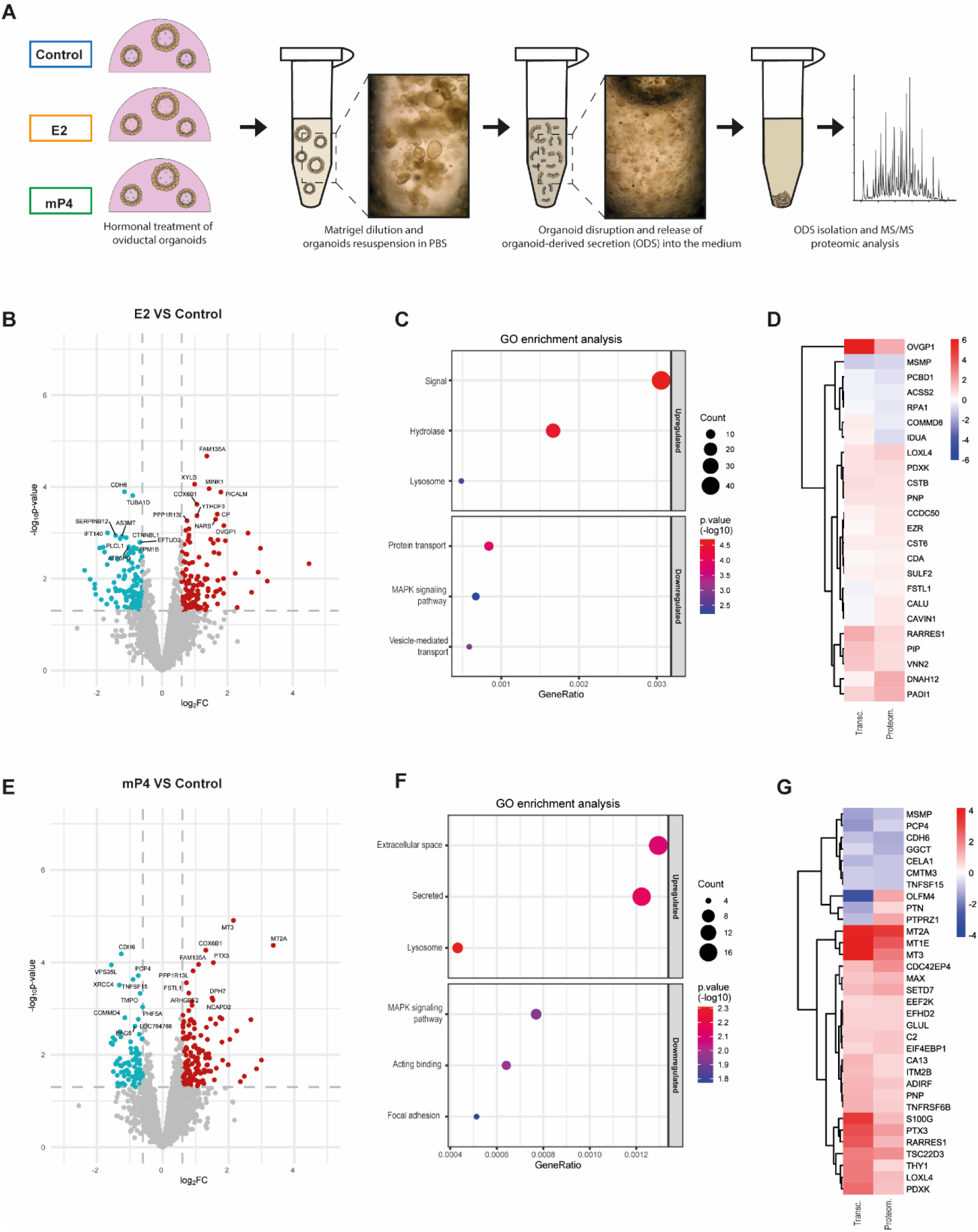
Proteomic characterization of organoid-derived secretion (ODS) following hormonal stimulation. (A) Schematic representation of the experimental design. Organoids from Control, E2, and mP4 groups were resuspended in PBS and disrupted to release ODS accumulated in the inner lumen, n = 3 independent biological replicates. Proteins released into the PBS were subsequently analysed by liquid chromatography–tandem mass spectrometry (LC-MS/MS). (B) Volcano plot of differentially expressed proteins in the E2 group vs. Control. (C) Dot plot of enrichment analysis of up- and downregulated proteins between E2 and Control groups. (D) Heatmap comparing differentially expressed genes in organoid transcriptomes (Transc.) with differentially expressed proteins in ODS proteomes (Proteom.) between E2 and Control groups. (E) Volcano plot of differentially expressed proteins in the mP4 group vs. Control. (F) Dot plot of enrichment analysis of up- and downregulated proteins between mP4 and Control groups. (G) Heatmap comparing top differentially expressed genes in organoid transcriptomes (Transc.) with differentially expressed proteins in ODS proteomes (Proteom.) between mP4 and Control groups.

Comparative analysis revealed that estradiol treatment upregulated 148 and downregulated 122 proteins relative to control, whereas progesterone altered 223 proteins (141 upregulated, 82 downregulated; Fig. 2B, E, Supplementary Table 5). PCA clearly separated the samples according to treatment, indicating distinct proteomic profiles (Supplementary Figure 2D). In the E2 group, enriched categories included signal peptides, hydrolases, and lysosomal proteins (Fig. 2C and Supplementary table 6), consistent with active secretory remodelling. Notably, classical oviductal fluid components such as *OVGP1*, *PIP*, *SELENOM*, and lysosomal enzymes (*HEXB*, *CTSB*) were upregulated, reflecting the physiological follicular-phase environment.

In mP4-treated ODS, GO enrichment analysis highlighted extracellular matrix proteins and protease regulators such as *CTSZ* and *TIMP1*, together with characteristic luteal-phase proteins (*HP*, *TGM2*) (Fig. 2F and Supplementary table 6). MAPK and cytoskeletal pathways were downregulated, suggesting reduced epithelial turnover and a shift toward a more quiescent, differentiated state. Integration of transcriptomic and proteomic datasets revealed strong concordance in the regulation of hormone-dependent secreted proteins (Fig. 2D, G), confirming that the observed secretory changes are transcriptionally driven.

### Multi-omics factor analysis identifies coherent hormonal signatures linking transcriptional and secretory regulation

To uncover coordinated molecular programs across transcriptomic and proteomic layers, we performed MOFA. Five latent factors explained 42.6% of transcriptomic and 82.0% of proteomic variance (Fig. 3A). Among them, Factor 2 strongly correlated with mP4 treatment (*p* = 0.004) and Factor 3 with E2 treatment (*p* = 0.027; Fig. 3C–D). Principal component projection of these factors clearly segregated samples by hormonal condition (Fig. 3B).

**Figure 3.**
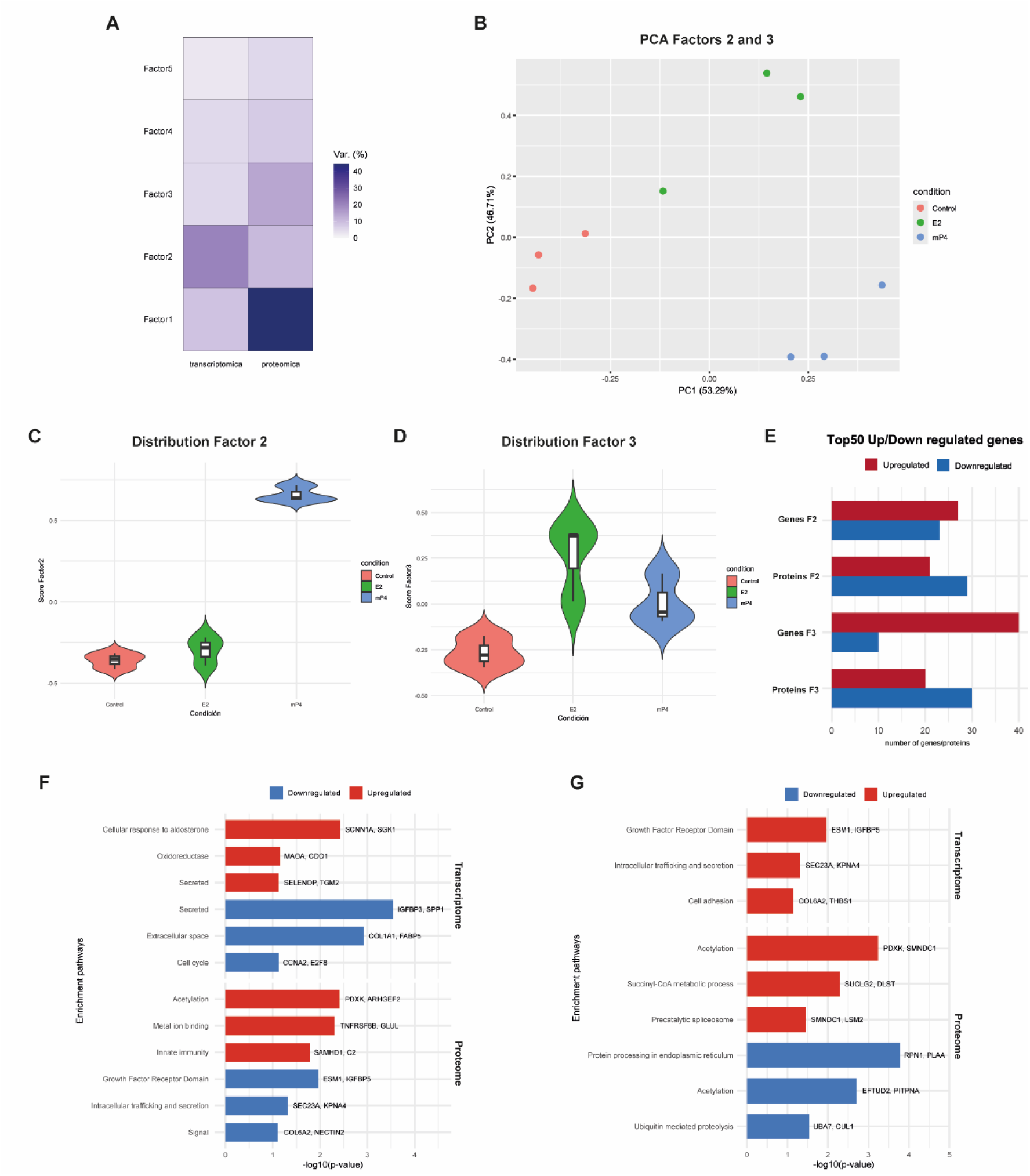
Integrated analysis of transcriptomic and proteomic datasets and hormone-dependent clustering. (A) Percentage of variance explained by each factor in the different omics views for the selected MOFA model. (B) Principal component analysis (PCA) after filtration of factors 2 and 3. (C-D) Violin plots representing the distribution of factors 2 and 3 according to hormonal treatment. (E) Bar graphic with top 50 genes and proteins up and downregulated for each factor. (F-G) Gene ontology enrichment of differentially expressed genes of Top 50 genes/proteins of factor 2 and factor 3 terms upregulated are shown in red, and downregulated terms in blue.

In the mP4-associated Factor 2, genes and proteins defining the progesterone response were enriched in extracellular remodelling and oxidative metabolism. *SELENOP*, *TGM2*, and *SGK1* were among the top contributors, reflecting the role of progesterone in redox and osmotic homeostasis (Fig. 3E, F). At the proteomic level, upregulated enzymes such as PDXK and GLUL indicated enhanced amino acid and vitamin metabolism, potentially supporting early embryo nutrition (Fig. 3E, F).

The E2-associated Factor 3 displayed a dominant contribution of genes related to growth factor signalling and secretion, including *ESM1*, *IGFBP4*, and *SEC23A*, as well as metabolic proteins supporting biosynthetic and mRNA processing activities (*SUCLG2*, *DLST*, *SMNDC1*, Fig. 3E, G). Together, MOFA integration revealed that hormone-specific transcriptional programs are mirrored at the secretory level, establishing the mechanistic coupling between epithelial differentiation and the composition of oviductal secretions.

### Estradiol-stimulated organoid secretions enhance sperm capacitation and acrosome reaction

To assess the functional relevance of ODS, we examined their capacity to modulate sperm physiology. Bull spermatozoa were incubated under capacitating conditions with ODS obtained from control, E2-, or mP4-treated organoids, with preovulatory oviductal fluid (BOF-LF) serving as a positive control and PBS as a negative control (Fig. 4A).

**Figure 4.**
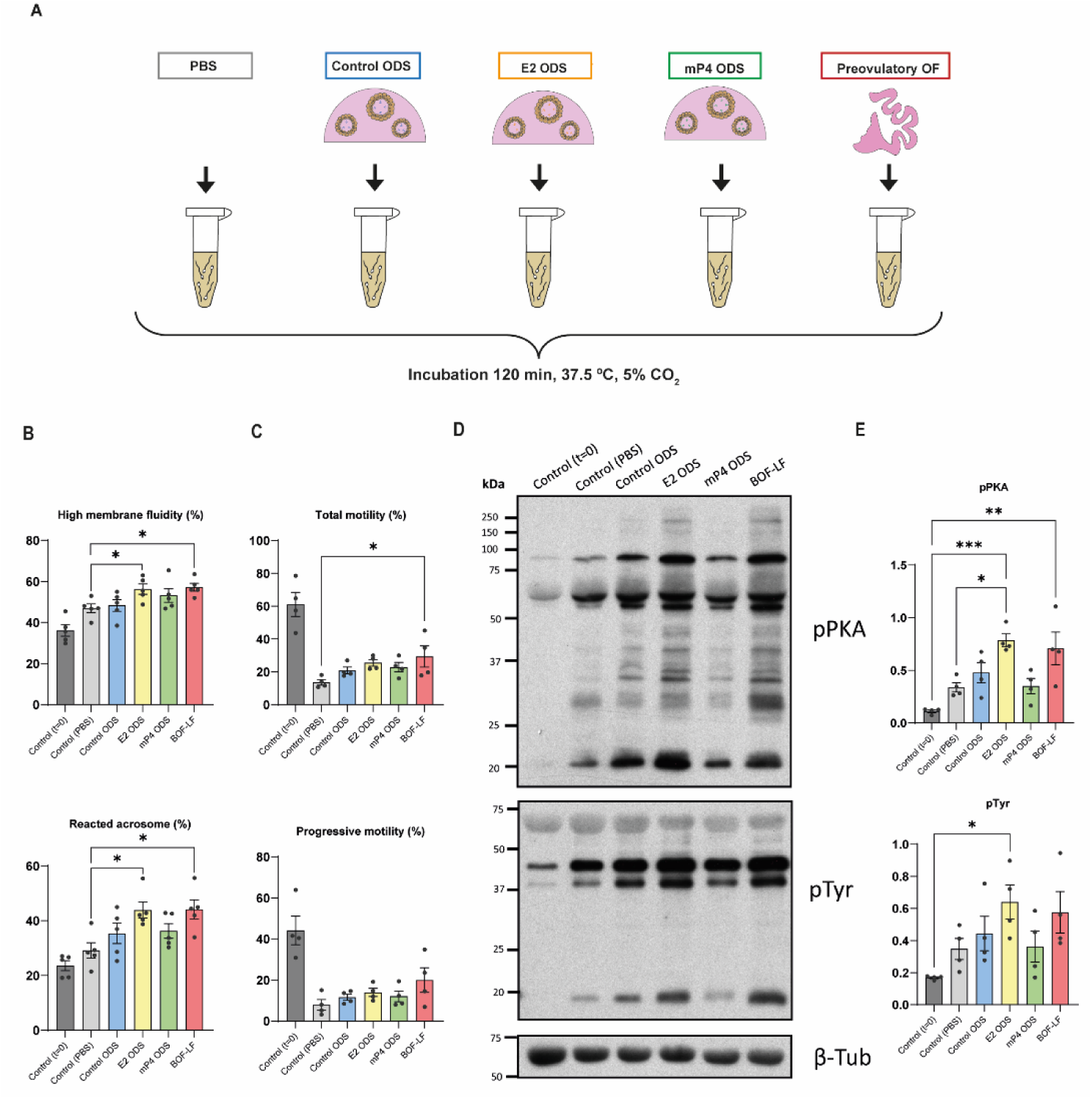
Functional analysis of organoid-derived secretions (ODS) in sperm capacitation. (A) Schematic representation of the experimental design. Bovine spermatozoa were incubated with ODS from different hormonal treatments (Control, E2, mP4), with preovulatory oviductal fluid (OF) as a positive control, and with PBS as a negative control. (B) Membrane status assessed as the percentage of spermatozoa with high membrane fluidity, and acrosome reaction evaluated by flow cytometry. (C) Motility parameters after two hours of incubation. Total and progressive motility are shown as bar plots. (D) Western blot analysis of pPKA substrates, pTyr, and β-tubulin as a loading control. (E) Quantification of Western blot signals for pPKA substrates and pTyr normalized to β-tubulin. For statistical analysis, differences were assessed using one-way ANOVA followed by Tukey’s multiple comparison test (p≤0.05). Data are presented as mean ± SEM. Statistical significance is indicated as *p < 0.05, **p < 0.01, ***p < 0.001.

After 2 hours of incubation, spermatozoa exposed to ODS-E2 displayed increased membrane fluidity and a significantly higher proportion of acrosome-reacted cells compared with controls (*p* < 0.05; Fig. 4B). Similarly, total and progressive sperm motility tended to increase with ODS-E2, with significant enhancement observed in the BOF-LF group, while other motility parameters remained unchanged (Fig. 4C and Supplementary table 7). Western blot analysis revealed enhanced phosphorylation of PKA substrates and tyrosine residues (pPKA and pTyr) (Fig. 4D–E), both molecular hallmarks of capacitation [31–33]. The capacitating effect of ODS-E2 paralleled that observed with native follicular-phase oviductal fluid, whereas ODS-mP4 had minimal effect, consistent with its association with the post-fertilization luteal phase [34].

Collectively, these data demonstrate that estradiol-stimulated ODS reproduce the functional activity of native oviductal fluid by promoting sperm capacitation and acrosome reaction. The organoid-derived secretome therefore captures not only the molecular but also the bioactive properties of the *in vivo* oviductal environment.

## Discussion

In this study, we established a hormonally responsive bovine oviductal organoid model that faithfully reproduces the molecular, structural, and functional features of the native oviduct. Through integrated transcriptomic and proteomic analyses, we demonstrated that these organoids recapitulate oestrous cycle–specific programs and generate bioactive secretions capable of modulating sperm physiology [8, 9, 34]. This work provides the first mechanistic evidence that organoid-derived secretions can mimic both the molecular complexity and biological activity of oviductal fluid, offering a physiologically relevant and sustainable platform to study gamete–maternal communication.

Our findings confirm that oviductal epithelial cells retain intrinsic endocrine responsiveness when cultured in a three-dimensional architecture. Estradiol promoted a secretory and metabolically active phenotype, while progesterone induced extracellular remodelling and epithelial quiescence, patterns that closely parallel the *in vivo* follicular and luteal phases. These responses are consistent with previous transcriptomic studies of the bovine oviduct and highlight the capacity of organoids to preserve hormonal sensitivity and transcriptional plasticity [30]. By using a defined hormonal regimen, we achieved precise temporal control of epithelial differentiation and secretion, overcoming the variability associated with *ex vivo* tissue or fluid collection. Our results also confirm that oviductal organoids are a suitable model for studying the oviductal secretome, due to their high proportion of secretory cells and their apical-in orientation, which allows the accumulation and subsequent isolation of components secreted by the organoids.

Integration of transcriptomic and proteomic datasets through multi-omics factor analysis revealed coherent regulation of endocrine-dependent programs, with estradiol and progesterone defining distinct epithelial states. Estradiol-induced upregulation of secretory and vesicle-trafficking genes and proteins, including ESM1, IGFBP4, and SEC23A, alongside luminal components such as OVGP1 and SELENOM, suggests that the organoids adopt a preovulatory, pro-secretory phenotype that actively modulates the oviductal microenvironment to support sperm–oocyte interactions [3]. Conversely, progesterone-dependent enrichment of extracellular matrix regulators, redox-related enzymes, and secreted proteins such as TGM2 and HP reflects a luteal-phase state favoring epithelial quiescence, matrix remodeling, and early embryonic support. The concordance between transcriptomic and proteomic signatures underscores the integrated regulation of these hormonally driven programs [2]. The strong concordance between gene expression and secreted protein profiles demonstrates that oviductal organoids maintain the integrated transcriptional and post-transcriptional regulation underlying physiological hormone responses.

Previous studies have demonstrated that oviductal fluid influences sperm capacitation [9, 35], oocyte maturation [36, 37] and embryo development [11, 38] across species, including bovine, porcine and human models [24, 39–42]. However, the variability and limited availability of post-mortem oviductal secretions restrict their experimental and clinical use. Co-culture with oviductal epithelial cells has also been shown to promote gamete interaction [43]. These observations provide a strong rationale for assessing the functional competence of ODS in supporting gamete and embryo development. Recent advances in organoid culture have introduced apical-out systems [44], which expose the apical surface directly to the external medium and are particularly valuable for studies of embryo-maternal interactions with a direct apical access. However, in the context of oviductal secretions, the apical-in orientation used in our study provides a unique advantage by allowing the accumulation of concentrated secreted factors within the lumen, facilitating proteomic profiling and functional assays such the effect of gamete or embryo *in vitro* culture.

Proteomic characterization of ODS further confirmed their molecular and functional fidelity to native oviductal fluid. ODS contained canonical oviductal proteins such as OVGP1, PIP, SELENOM, HP*, and* TGM2, whose abundance changed according to hormonal treatment in agreement with *in vivo* profiles [30, 45–50]. The enrichment of OVGP1, NTS, and CD9 in E2-treated ODS provides a mechanistic basis for the enhanced sperm capacitation and acrosome reaction observed, linking molecular composition directly to functional outcomes [8, 9, 34]. Similarly, luteal-phase proteins identified in mP4 ODS, such as TGM2 and HP, reflect the role of organoids in maintaining an extracellular milieu conducive to early embryo support, consistent with in vivo oviductal physiology [45, 47]. This indicates that organoid-derived secretions not only replicate the molecular composition of the oviductal environment but also recapitulate its physiological role in modulating gamete function. The enrichment of *OVGP1* and *NTS* under estradiol stimulation, two proteins previously shown to enhance sperm–zona pellucida interaction and capacitation signalling, likely contributes to these effects [51–53]. Such findings bridge molecular identity and biological function, demonstrating that ODS represent a bioactive surrogate of the native oviductal milieu.

Beyond their biological significance, these results have broad methodological and translational implications. Oviductal organoids provide a renewable, controllable, and ethical alternative to animal-derived reproductive fluids, enabling systematic exploration of hormonal regulation, secretion, and gamete interaction under defined conditions. The capacity to generate physiologically relevant secretions *in vitro* opens new possibilities for improving assisted reproductive technologies by supplementing gamete or embryo culture media with organoid-derived factors that restore essential reproductive cues absent in conventional formulations. Moreover, this system can be adapted to investigate oviductal pathophysiology, environmental influences on fertility, and interspecies differences in reproductive strategies.

While this model reproduces major aspects of oviductal physiology, several refinements could further enhance its utility. The cellular composition of the organoids, particularly the ratio of ciliated to secretory cells, remains to be quantified, and integrating microfluidic approaches could reproduce luminal flow and mechanical cues characteristic of the native oviduct. Standardization of donor material and long-term culture stability will also be important for translational reproducibility. Future studies should aim to identify the specific molecular mediators within ODS responsible for modulating sperm and embryo physiology, combining proteomics with lipidomic and extracellular vesicle profiling. Ultimately, coupling oviductal organoids with gamete and embryo co-cultures in dynamic environments may yield a next-generation platform to dissect fertilization and early development under conditions that faithfully reproduce *in vivo* physiology.

In summary, our findings establish bovine oviductal epithelial organoids as a robust and mechanistically informative model that mirrors the hormonal regulation and secretory landscape of the oviduct. The ability of their secretions to promote sperm capacitation demonstrates functional equivalence to native oviductal fluid and underscores their potential as a biomimetic tool for reproductive biology. This work not only advances our understanding of oviductal epithelial dynamics but also provides a foundation for developing physiologically inspired strategies to improve the efficiency and fidelity of assisted reproductive technologies.

## Acknowledgments

Thank you for support from the Advanced Light Microscopy Facility (SMOA) at the Centro de Biología Molecular Severo Ochoa (Madrid, Spain). Special thanks to Dr. Vasiliki Lalioti for her help with fluorescence microscopy.

## Authors’ contributions

SNS, JRA, PC, and VPG contributed to the conception and design of the study. Material preparation and experimental analyses were performed by SNS, JRA, NHD, and AFM. Data processing, bioinformatic, and statistical analyses were performed by SNS. The first draft of the manuscript was written by SNS and VPG. All authors commented on previous versions of the manuscript and approved the final version.

## Funding

This work was supported by Grant PLEC2022-009246 funded by MCIN/AEI/10.13039/501100011033 and by the “European Union NextGenerationEU/PRTR”.

## Data availability

All data generated or analysed during this study are included in this published article and its supplementary information files. Genome-wide sequencing data have been deposited in the European Nucleotide Archive (ENA) under accession number PRJEB104075, and the mass spectrometry proteomics data have been deposited to the ProteomeXchange Consortium via the PRIDE partner repository with the dataset identifier PXD071140.

## Conflict of interest

The authors declare that they have no conflict of interest.

---

**Supplementary Figure 1.**
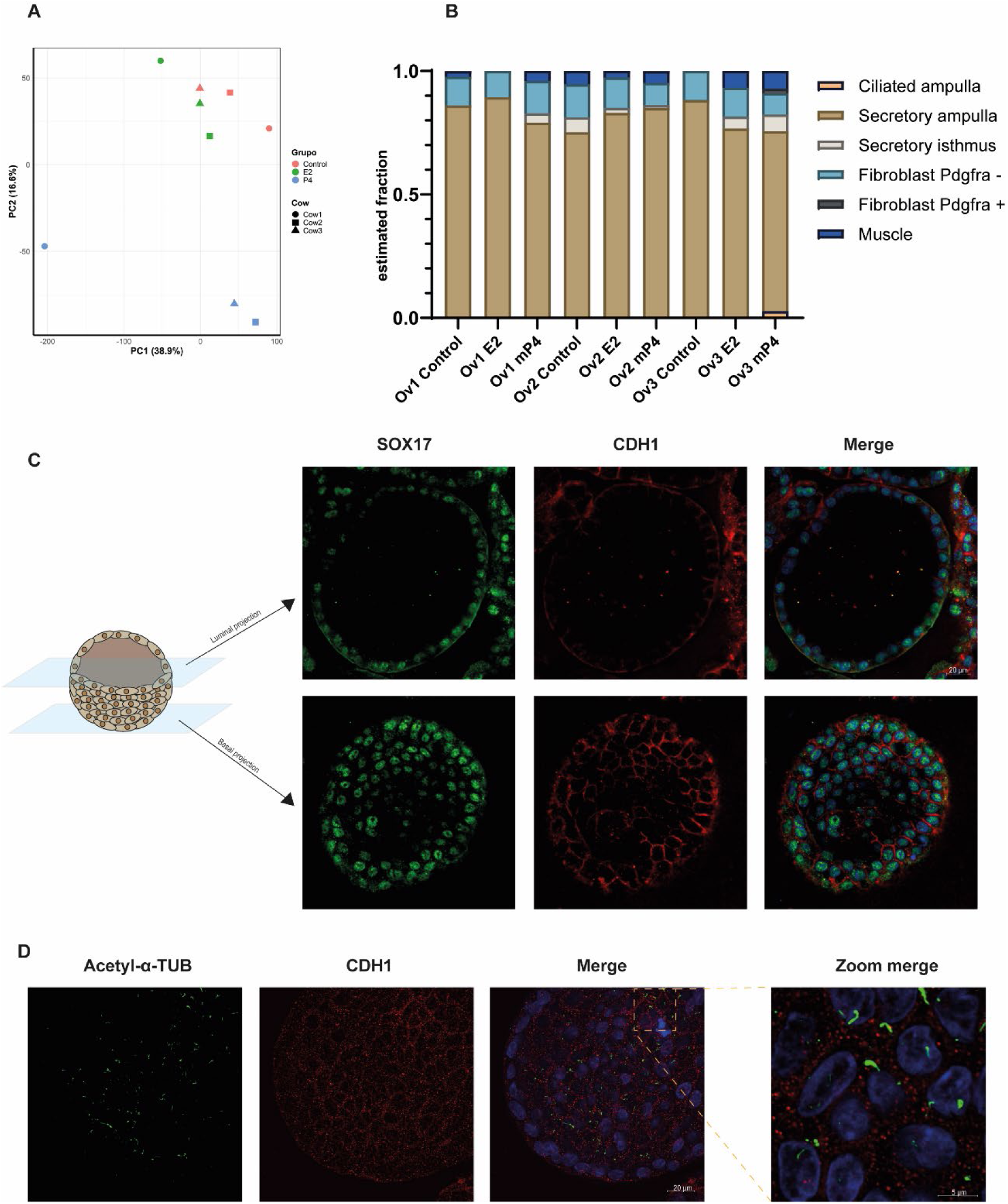
(A) Principal component analysis (PCA) of bulk RNA-seq data from control organoids and organoids treated with estradiol or mP4. (B) Bar graph showing the estimated fractions of cell types in bovine oviductal organoids after deconvolution using single-cell RNA-seq data. (C) Immunofluorescence staining of oviductal organoids showing basal and luminal projections, stained for SOX17 (green, secretory marker), CDH1 (red), and DAPI (blue). (D) Immunofluorescence staining of oviductal organoids stained for acetyl-α-tubulin (green), CDH1 (red), and DAPI (blue).

**Supplementary Figure 2.**
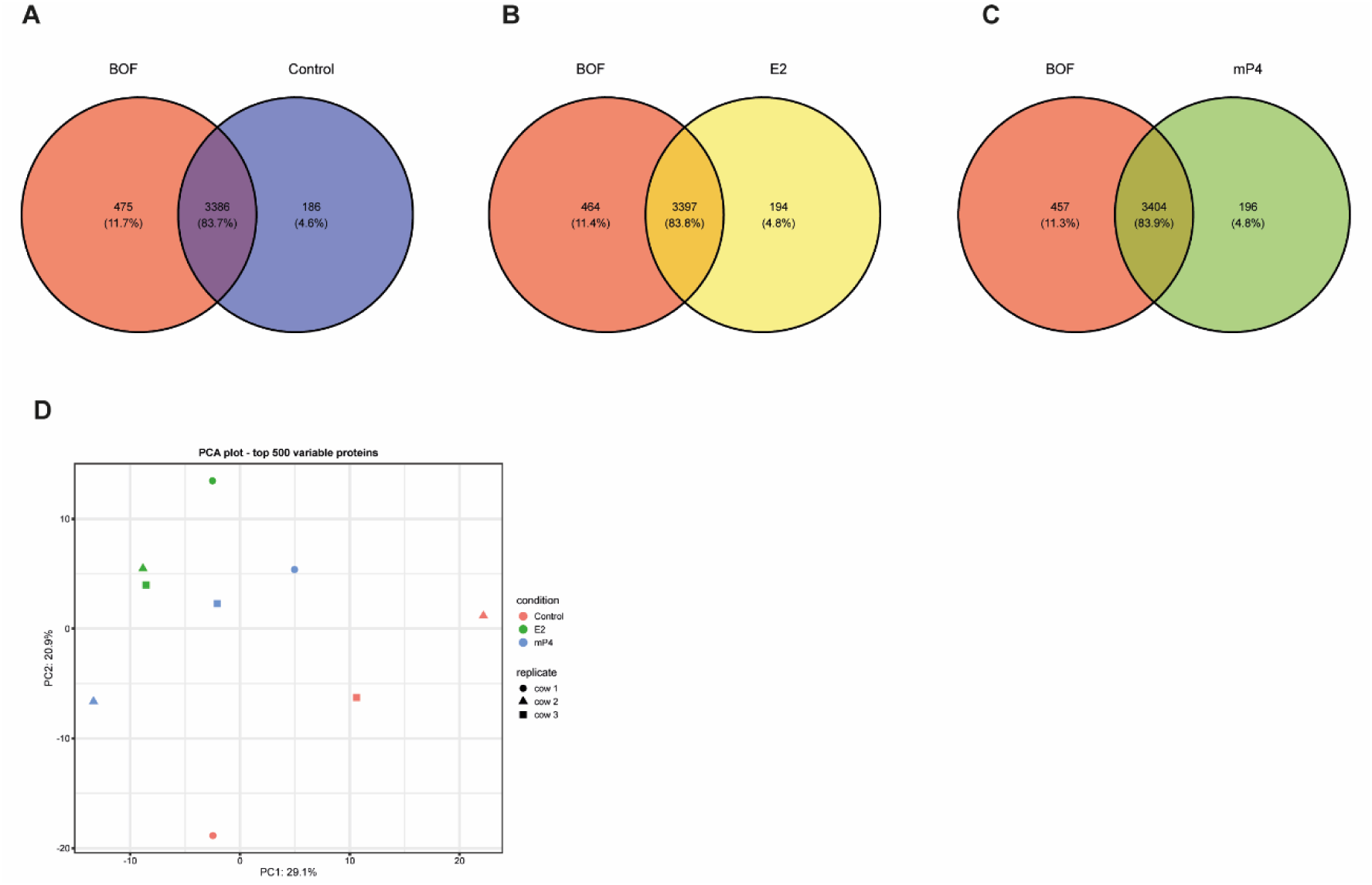
(A–C) Venn diagrams showing proteins identified in bovine oviductal fluid from the early luteal phase (BOF) compared with organoid-derived secretions (ODS) from control organoids (A), estradiol-treated organoids (E2, B), and organoids treated with estradiol, medroxyprogesterone, and cAMP (mP4, C). (D) Principal component analysis (PCA) of the proteomic profiles of ODS from the three hormonal treatments.

## Notes

### Competing Interest Statement

The authors have declared no competing interest.

